# Sleep induced by mechanosensory stimulation provides cognitive and health benefits in *Drosophila*

**DOI:** 10.1101/2024.07.10.602891

**Authors:** Sho Inami, Kyunghee Koh

## Abstract

**Study Objectives:** Sleep is a complex phenomenon regulated by various factors, including sensory input. Anecdotal observations have suggested that gentle rocking helps babies fall asleep, and experimental studies have verified that rocking promotes sleep in both humans and mice. Recent studies have expanded this understanding, demonstrating that gentle vibration also induces sleep in *Drosophila*. Natural sleep serves multiple functions, including learning and memory, synaptic downscaling, and clearance of harmful substances associated with neurodegenerative diseases. Here, we investigated whether vibration-induced sleep provides similar cognitive and health benefits in *Drosophila*.

**Methods:** We administered gentle vibration to flies that slept very little due to a forced activation of wake-promoting neurons and investigated how the vibration influenced learning and memory in the courtship conditioning paradigm. Additionally, we examined the effects of VIS on synaptic downscaling by counting synapse numbers of select neurons. Finally, we determined whether vibration could induce sleep in *Drosophila* models of Alzheimer’s disease (AD) and promote the clearance of Amyloid β (Aβ) and Tubulin Associated Unit (TAU).

**Results:** Vibration-induced sleep enhanced performance in a courtship conditioning paradigm and reduced the number of synapses in select neurons. Moreover, vibration improved sleep in *Drosophila* models of AD, promoting the clearance of Aβ and TAU.

**Conclusions:** Mechanosensory stimulation offers a promising non-invasive avenue for enhancing sleep, potentially providing associated cognitive and health benefits.

**Significance Statement:** Sleep is critical for a healthy mind and body, and sleep disturbances are commonly associated with neurodegenerative diseases such as Alzheimer’s disease. Sleep is influenced by sensory input, and mechanical stimulation, such as gentle rocking and vibration, has been shown to promote sleep in various species, including humans, mice, and fruit flies. This study demonstrates that gentle vibration not only helps flies sleep better but also improves their performance in a learning and memory task and makes their brains more efficient in clearing harmful substances. Notably, vibration can facilitate the clearance of Amyloid β and the TAU proteins, which accumulate in Alzheimer’s disease. These results highlight the potential for gentle mechanosensory stimulation to promote sleep and cognitive health.

## Introduction

Sleep is a complex process regulated by many factors, including sensory input.^1^ Anecdotal observations have long suggested the soothing effect of gentle rocking on infants, leading to experimental studies confirming its sleep-promoting effects in both infants and adults as well as mice.^2–5^ Furthermore, rocking-induced sleep has been shown to enhance memory in adult humans.^6^ We and others have recently reported that gentle mechanical stimulation induces sleep in *Drosophila* as well.^7,8^ The studies found that various types of mechanosensory stimuli, including vibrations and orbital motions, promoted sleep in male and female flies across multiple control genetic backgrounds. These findings provide an opportunity to investigate the benefits of sleep induced by mechanosensory stimuli using the *Drosophila* model system.

Sleep serves multiple functions, including learning and memory, synaptic plasticity, and clearance of harmful metabolites.^9,10^ Substantial evidence supports a conserved role for sleep in enhancing learning and memory.^11–14^ Sleep deprivation negatively impacts cognitive performance across species, including humans, rodents, and flies.^15–18^ Conversely, inducing sleep by activating sleep-promoting neurons or administering drugs restores performance in animals with impaired learning and memory.^19–23^ Improved learning and memory during sleep may be mediated by the different ways synaptic connections are altered during sleep and wakefulness. The “synaptic homeostasis” hypothesis proposes that there is an overall increase in synaptic strength while awake, while there is an overall decrease in synaptic connections during sleep. ^24,25^ Consistent with the hypothesis, a number of studies in mice, zebrafish, and flies have demonstrated that synapses are generally weakened during sleep, and waking activities lead to synaptic strengthening and a widespread increase in synaptic proteins.^26–31^

Another critical role of sleep is in enhancing the activity of the glymphatic system in the brain and thus promoting the clearance of toxic substances, including Amyloid β (Aβ) and Tubulin Associated Unit (TAU) linked to Alzheimer’s disease (AD).^32–34^ Sleep disorders are a common comorbidity in Alzheimer’s disease AD patients.^35–37^ Accumulating evidence suggests that the relationship between sleep and AD is bi-directional; AD causes sleep disturbances, while disrupted sleep accelerates AD pathologies.^38,39^ Disrupted sleep may contribute to cognitive deficits in AD patients through defective metabolite clearance.^40,41^ Importantly, improved sleep can ameliorate memory deficits in a *Drosophila* model of AD,^42^ suggesting that sleep-related interventions could be an effective strategy for treating AD patients. Since sleep disruptions occur early in the course of AD,^36,43^ improving sleep may be a particularly effective strategy for slowing down the progression of AD. However, whether vibration can promote sleep in AD patients or animal models of AD has not been investigated.

We previously showed that vibration-induced sleep (VIS) in *Drosophila* exhibits a few characteristics of spontaneous sleep. VIS is associated with reduced arousability, a hallmark of sleep, and flies exhibit decreased sleep after vibration, suggesting sleep credit accumulates during VIS. In this study, we further investigated the parallels between VIS and spontaneous sleep, focusing on the cognitive and health benefits. Our results show that VIS enhances cognitive performance and facilitates synaptic downscaling. Moreover, our findings present evidence that vibration can enhance sleep in *Drosophila* models of Alzheimer’s disease, promoting the clearance of Aβ and TAU implicated in the condition. Gentle mechanical stimulation may offer a promising non-pharmacological avenue for enhancing sleep, potentially providing associated health and cognitive benefits, in individuals with sleep disorders and neurodegenerative diseases.

## Methods

### Fly stocks

Flies were raised on standard *Drosophila* medium containing cornmeal and molasses under a 12 h:12 h light/dark (LD) cycle. We obtained *Pdf*-*Gal4* (#80939), *R11H05*-*Gal4* (#45016), UAS-TDAG51::GFP (*tdGFP*, #35839), UAS-*dTrpA1* (#26263), UAS-*Syt1::GFP* (#6925), UAS-*Tau.2N4R* (#93610), and the background control strain iso31 (*w*^1118^, #3605) from the Bloomington *Drosophila* Stock Center; *R11H05*-*LexA* from the Janelia Fly Facility; LexAop-*TrpA1* and UAS-Aβ42.Arctic from Mark Wu (Johns Hopkins University); and *Da*-*GS* from Amita Sehgal (University of Pennsylvania). All fly lines were outcrossed to the iso31 control strain for five or more generations.

### Sleep analysis

Approximately 16 males and 16 females were housed together under a 12h:12h LD cycle until individual 3- to 5-day-old flies were loaded into glass tubes containing 5% sucrose and 2% agar. Experiments were performed at 25℃ except for those involving *TrpA1*. In those experiments, flies were raised at 22℃, monitored for 1 day at 22℃ to establish baseline sleep, and then monitored for 1 day at 28℃ or 30℃ to determine the effects of TrpA1 activation on sleep. The *Drosophila* Activity Monitoring (DAM) System (Trikinetics) was employed to record infrared beam breaks in 1-min bins. Sleep was defined as a period of no beam breaks lasting 5 min or more^44^ and analyzed using a MATLAB-based software SleepLab.^45^

### Generation of vibration stimuli

The experimental setup employed a multitube vortexer to generate complex vibratory stimuli, as described previously.^7^ Briefly, DAM monitors were placed approximately 40 cm above a multitube vortexer (Fisher Scientific) placed inside a temperature- and light-controlled incubator (VWR International). The vortexer intensity was set to 3, and the LC4 Light Controller (Trikinetics) was employed to control the timing and duration of the mechanical stimulation.

### Courtship conditioning

The courtship conditioning assay was performed as previously described.^46,47^ Briefly, a 3- to 5-day-old virgin male was placed with a 4 to 7-day-old mated female in a DAM monitor tube (4 mm in diameter, 5 cm long) for 1 h with (conditioned) or without (naive) a pre-mated female. Subsequently, each male fly was kept in isolation for 1 h until its courtship behavior toward a freeze-killed virgin female was assayed in a test chamber. The male courtship activity was evaluated manually to determine a courtship index (CI), the percentage of time spent performing courtship behaviors during a 10-min observation period or until copulation. Only male flies that showed courtship behaviors toward the mated female but did not copulate during conditioning were included.

### RU486 treatment

To induce expression of Aβ and TAU using *Da*-*GS*, flies were fed 500 μM RU486 (Sigma Aldrich). RU486, diluted in ethanol, was added to a standard *Drosophila* medium, and ethanol was used as the vehicle control. To minimize the degradation of RU486, flies were kept under constant dark (DD), and fresh food was provided every 3 days. For sleep measurements, flies were loaded into monitor tubes containing RU486 or ethanol diluted in 5% sucrose and 2% agar on Day 10. For immunohistochemistry experiments, flies were kept in vials containing a standard medium with RU486 or ethanol for 14 days.

### Immunohistochemistry

To perform whole-mount immunostaining, female brains were dissected and fixed in 4% paraformaldehyde (EMS) for 45 min at room temperature (RT). The brains were then washed three times in PBT (PBS with 0.3% Triton-X) and blocked in 1% normal goat serum in PBT for 1 h at RT. They were subsequently incubated with primary antibody at 4°C overnight. Following three washes in PBT, the brains were incubated with secondary antibodies at 4°C overnight. Rabbit anti-GFP (Molecular Probes), Alexa Fluor 488 goat anti-rabbit (Thermo Fisher Scientific), and Cy5 goat anti-mouse (Thermo Fisher Scientific) were used at 1:1000; mouse anti-Aβ (Covance) and mouse anti-TAU (DSHB) at 1:500. A Leica SP8 confocal microscope (Leica Microsystems) was employed to collect Z-sections at 1 μm intervals.

### Quantification of confocal images

We used Fiji (https://fiji.sc) to analyze confocal images. For quantification of synapse numbers, synapses were visualized using Syt1::GFP. A 3D image of the optic lobe was created from sequential confocal slices. The number of synapses for each optic lobe was counted using the spot detection algorithm in Fiji, with the diameter of the spot set to 0.5-1.0 μm, and the average number of synapses per optic lobe was calculated for each brain. To analyze Aβ and TAU accumulation, average projection images were used to quantify signal intensities of the entire brain.

### Statistical analysis

We performed statistical tests using Prism 10 (GraphPad). We determined whether the data were normally distributed using Kolmogorov–Smirnov tests. For a given data type, we used non-parametric tests for all comparisons if there was any group that deviated significantly from the normal distribution. For instance, although most sleep data were normally distributed, since there were a few that were not, we employed non-parametric tests for all sleep data. To compare pairs of groups, we performed unpaired Student’s *t*-tests for normal distributions and Mann–Whitney *U* tests for non-normal distributions. To compare 3 or more groups, one-way ANOVAs followed by Holm-Šídák’s *post hoc* tests were used for normally distributed data. If the groups were not normally distributed, Kruskal–Wallis tests followed by Dunn’s multiple comparisons tests were carried out.

## Results

### VIS rescues learning and memory deficits caused by sleep loss

To investigate whether VIS can rescue learning and memory defects caused by sleep loss, we first sought to determine whether vibration can induce sleep in flies that lost most of their sleep due to the activation of wake-promoting neurons.^48^ We employed the *Gal4*/UAS system to express the temperature-sensitive TrpA1 channel^49^ in wake-promoting neurons labeled by R11H05-*Gal4* (R11H05 neurons).^48^ We raised flies and monitored their baseline sleep at 22°C, a temperature at which TrpA1 channels are closed. TrpA1 channels open at temperatures above 25°C, triggering neural activation. Since activation of these wake-promoting neurons has a greater effect on nighttime sleep compared to daytime sleep,^48^ we raised the temperature at night. Without vibration, raising the temperature from 22°C to 28°C led to a dramatic reduction in sleep in flies expressing *TrpA1* (R11H05 > *TrpA1*) compared to control flies expressing *tdGFP* (R11H05 > *tdGFP*) (Figure 1, A-B). To administer gentle vibration, we placed *Drosophila* activity monitors on a shelf approximately 40 cm above a multi-tube vortexer.^7^ Consistent with our previous findings, we did not observe nighttime VIS in control flies,^7^ which likely reflects a ceiling effect since they had a high baseline sleep at night (Figure 1, A-B). In contrast, gentle vibration resulted in a partial but significant recovery of the sleep lost due to R11H05 neuron activation. These findings allowed us to examine whether VIS can rescue memory deficits caused by sleep loss.

**Figure 1.**
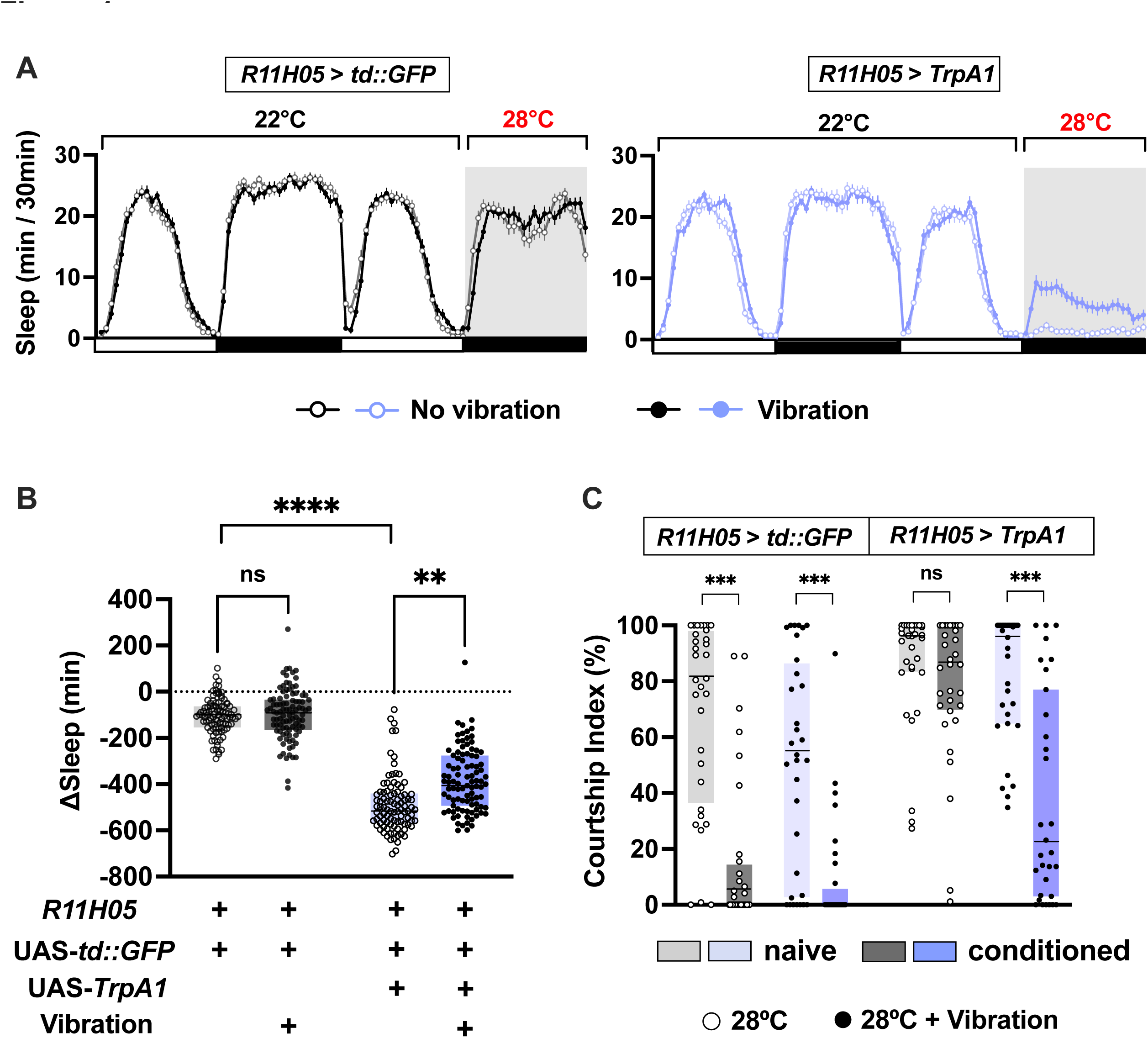
VIS alleviates learning and memory impairments caused by sleep loss. **(A)** Sleep patterns of male flies expressing *TrpA1* in wake-promoting neurons (R11H05 > *TrpA1*) and control flies expressing membrane-localized *tdGFP* (R11H05 > *tdGFP*). We kept the flies at 22°C for baseline sleep measurements and raised the temperature to 28°C to activate the warmth-sensitive TrpA1 channel, as indicated above the brackets. One group of flies received continuous vibration for 12 h starting at Zeitgeber time (ZT) 12 (marked by the gray areas), while the other group served as “no-vibration” controls. Mean and SEM are shown. N = 94-96. **(B)** Sleep changes (nighttime sleep at 28°C vs. 22°C) in the flies shown in Figure 1A. **(C)** Courtship index of naive and conditioned males of the indicated genotypes with or without vibration. N = 31-32. The line within the box denotes the median, while the box illustrates the 25^th^ to 75^th^ percentiles. In all figures, ns, not significant, \**p* < 0.05, \*\**p* < 0.01, \*\*\**p* < 0.001, **** *p* < 0.0001. Kruskal–Wallis test followed by Dunn’s multiple comparisons test (B-C).

To determine the effects of VIS on learning and memory, we employed the well-established, ecologically-relevant courtship conditioning assay.^46,47,50,51^ In this assay, male flies are “conditioned” to suppress courtship through repeated rejections by non-receptive mated females, and memory is measured by the degree of courtship suppression toward virgin females. Conditioning for 1 h can generate memory that persists for several hours in wild-type flies.^47,51^ We performed courtship conditioning on the day following sleep manipulations by vibration and activation of waking-promoting neurons, as described above. Control flies expressing *tdGFP* (R11H05 > *tdGFP*) exhibited reduced courtship after conditioning, regardless of vibration (Figure 1, C). In contrast, in the absence of vibration, flies that slept very little due to activated R11H05 neurons (R11H05 > *TrpA1*) exhibited little or no courtship suppression. This finding confirms previous results showing that sleep loss negatively impacts performance in courtship conditioning.^20,28^ Importantly, in the presence of vibration, flies with activated R11H05 neurons showed robust courtship suppression (Figure 1, C). Taken together, these results show that vibration can restore cognitive performance as well as sleep in flies with severely reduced sleep due to activated wake-promoting neurons.

### VIS rescues synaptic downscaling deficits caused by sleep loss

Previous studies have shown that a group of circadian neurons expressing pigment-dispersing factor (PDF), called ventral lateral neurons (LN_v_s),^52^ undergo synaptic downscaling during sleep.^28,30^ To determine whether synaptic downscaling also occurs during VIS, we expressed GFP-tagged presynaptic protein, Synaptotagmin 1 (Syt1)^53^ in LN_v_s using *Pdf*-Gal4 and examined synaptic terminals of large LN_v_s (l-LN_v_s), a subset that sends projections to the optic lobes. We again employed TrpA1-mediated thermogenetic activation of R11H05 neurons to induce sleep loss. Since we used the *Gal4*/UAS system to express *Syt1::GFP* in PDF-expressing neurons, we employed an orthogonal expression system, *LexA*/LexAOp, to express *TrpA1* in R11H05 neurons.^54^ The *LexA*/LexAOp system typically leads to a lower level of expression than the *Gal4*/UAS system, and thus we used a higher temperature than in the experiments employing R11H05-*Gal4* to activate TrpA1. Raising the temperature to 30°C caused a severe loss of nighttime sleep in flies expressing *TrpA1* in R11H05 neurons (*Pdf* > *Syt1::GFP*, R11H05 > TrpA1) (Figure 2, A-B). In contrast, control flies not carrying LexAop*-TrpA1* (*Pdf* > *Syt1::GFP*, R11H05 > +) showed only a modest sleep reduction upon temperature increase. The number of the presynaptic puncta in the optic lobe marked by Syt1::GFP was greater in sleep-deprived flies relative to control flies (Figure 2, C-D), confirming previous findings that sleep deprivation resulted in an increased number of synaptic terminals.^28,30^ To test whether VIS downscales synaptic connections like natural sleep, we applied gentle vibration at night during thermogenetic sleep deprivation. Vibration significantly rescued the sleep loss due to the activation of R11H05 neurons (Figure 2, A-B). Importantly, vibration fully restored the number of presynaptic terminals in flies with activated R11H05 neurons (Figure 2, C-D), supporting the hypothesis that VIS, like natural sleep, promotes downscaling of synapses.

**Figure 2.**
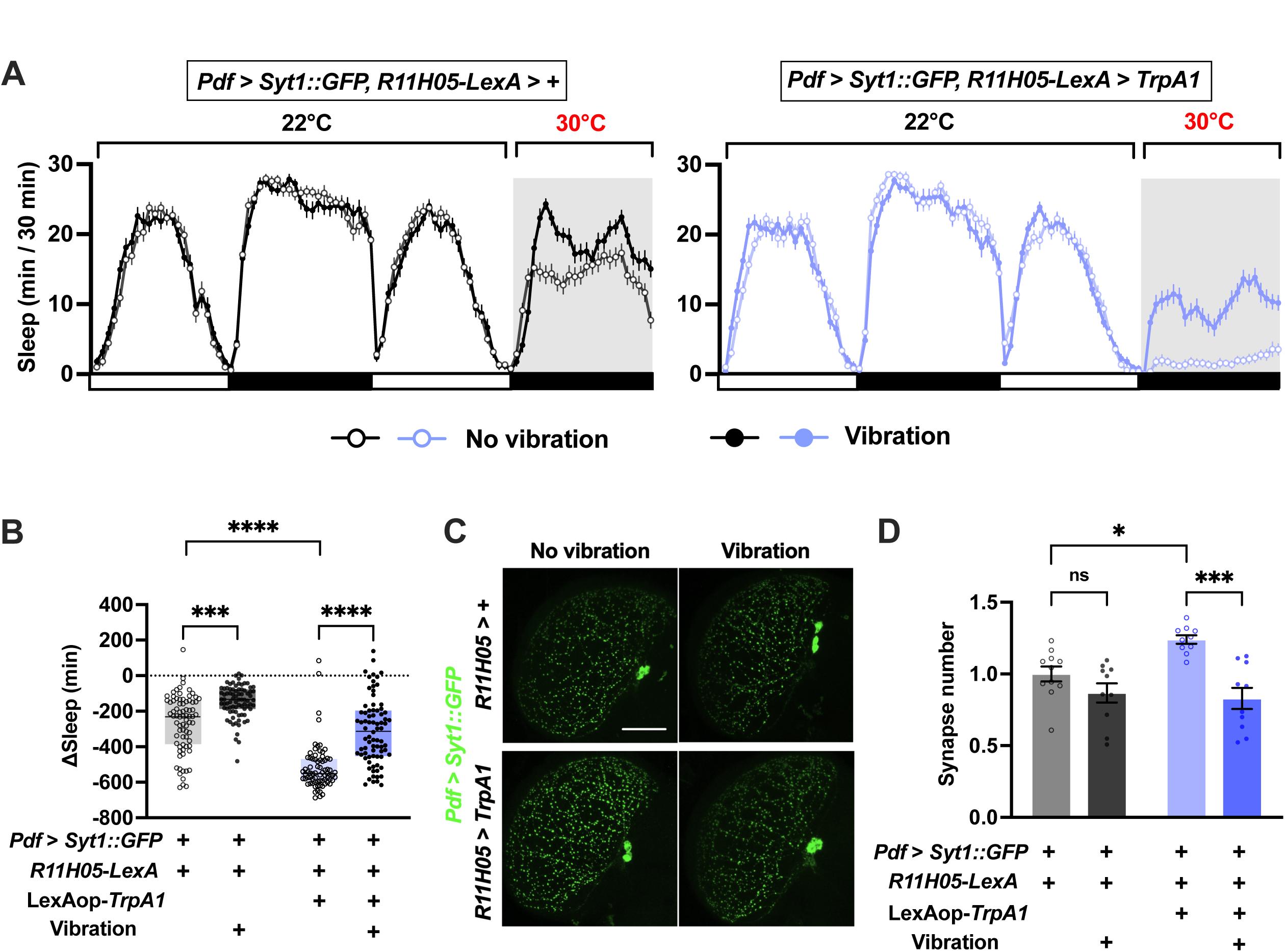
VIS counteracts the synaptic downscaling defects caused by sleep loss. **(A)** Sleep patterns of male flies from the specified genotypes. Flies were kept at 22°C for baseline sleep measurements, and the temperature was raised to 30°C to activate TrpA1, as indicated above the brackets. The gray area indicates 12-h vibration starting at ZT 12. Mean and SEM are shown. N = 79-80. **(B)** Sleep changes (nighttime sleep at 30°C vs. 22°C) in the flies shown in Figure 2A. The line within the box denotes the median, while the box illustrates the 25^th^ to 75^th^ percentiles. **(C)** Representative confocal images of the synaptic puncta of l-LNvs in the optic lobe. Synapses were marked by Syt1::GFP expression under the control of *Pdf*-*Gal4*. TrpA1 was expressed using R11H05-*LexA*, and the temperature was raised from 22°C to 30°C for 12 h starting at ZT12 to activate the wake-promoting R11H05 neurons in the presence or absence of vibration. Flies lacking LexAop-*TrpA1* served as genotypic controls. The brain was dissected between ZT0 and ZT1. Scale bars represent 50μm. **(D)** Quantification of the number of synapses in the optic lobes of the indicated genotypes and conditions. Error bars indicate SEM. N=10-11 brains. Kruskal–Wallis test followed by Dunn’s multiple comparisons test (B); one-way ANOVA followed by Holm-Šídák’s multiple comparisons test (D).

### VIS ameliorates Aβ and TAU accumulation in *Drosophila* models of AD

Our previous study showed that vibration does not induce sleep in all short-sleeping mutants.^7^ We, therefore, investigated whether vibration can induce sleep in a *Drosophila* model of AD. We employed an AD model expressing Aβ42.Arctic, an aggregation-prone mutant form of Aβ, which was shown to exhibit reduced sleep.^55^ We ubiquitously expressed Aβ42.Arctic in the adult using the *daughterless*-Geneswitch (*da*-GS), which contains an RU486-responsive conditional *Gal4* protein that allows temporal control of transgene expression.^56^ We fed adult flies with food containing 500 μM RU486 for 14 days and measured sleep during the last 3 days with or without vibration. Consistent with previous results,^55^ Aβ42.Arctic expression significantly reduced sleep amount in the no-vibration condition (Figure 3A-C). Importantly, sleep was markedly enhanced by vibration in flies expressing Aβ42.Arctic, showing that vibration is an effective way to promote sleep in a *Drosophila* model of AD.

**Figure 3.**
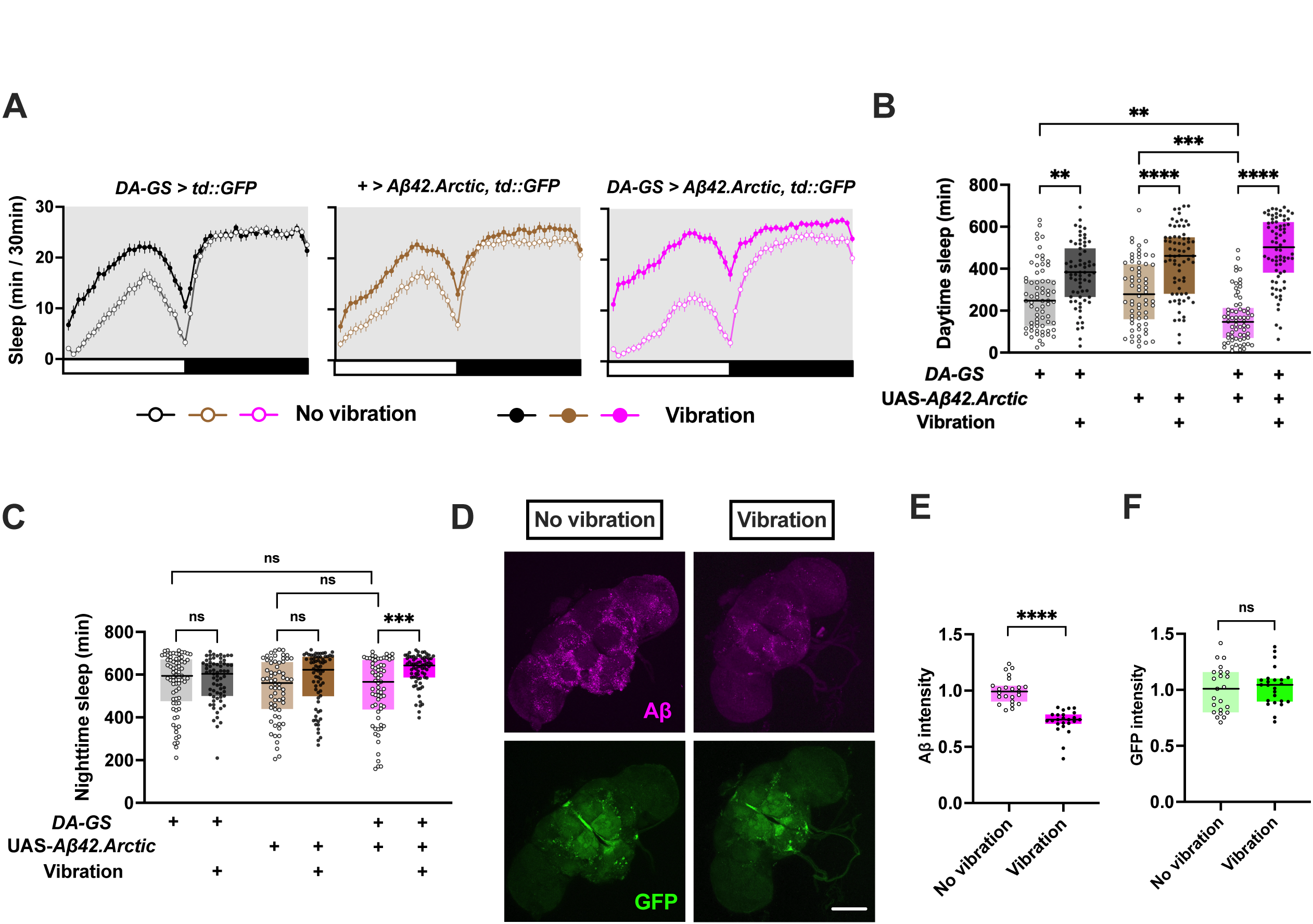
Vibration induces sleep in Aβ-expressing flies and reduces Aβ accumulation. **(A)** Sleep patterns of female flies from the specified genotypes and vibration conditions. All flies carried the UAS-*tdGFP* construct to confirm that Aβ42.Arctic expression did not cause non-specific effects on protein expression. Flies were exposed to continuous vibration for 3 days starting at ZT 0. Sleep patterns remained consistent over 3 days, and the sleep amount for each 30-min bin was averaged over the 3-day period. Mean and SEM are shown. N = 56-75. **(B-C)** Daytime (B) and nighttime (C) sleep for data shown in A. **(D)** Representative confocal images of immunolabeled Aβ and GFP signals in the whole brains of flies expressing Aβ.Arctic and tdGFP under the control of *da-GS*. Flies were fed 500 μM RU486 for 14 days before the brain dissection, and the vibration group received continuous vibration during the final 3 days. Scale bar: 50 μm. **(E)** Quantification of Aβ signal intensity normalized to the average intensity of the no-vibration condition. N = 23-24. **(F)** Quantification of GFP signal intensity normalized to the average intensity of the no-vibration condition. N = 23-24. The line within the box denotes the median, while the boxes illustrate the 25^th^ to 75^th^ percentiles (B, C, E). Kruskal–Wallis test followed by Dunn’s multiple comparisons test (B, C); Mann-Whitney *U* test (E, F).

Considering previous findings demonstrating that pharmacological and genetic induction of sleep in *Drosophila* models of AD can ameliorate Aβ accumulation,^42,55^ we tested whether VIS reduced Aβ accumulation. We employed flies expressing Aβ42.Arctic ubiquitously under the control of *da-*GS and fed 500 μM RU486 for 14 days. During the final 3 days, one group of flies received vibration while the control group did not. We compared the accumulation of Aβ in the brain using whole-mount immunostaining. Brains after 3 days of vibration showed a significant reduction of Aβ, compared to no-vibration control flies (Figure 3, D-E). GFP expression was not altered by vibration, demonstrating a specific effect on Aβ accumulation (Figure 3, D, F). These results provide evidence that VIS helps to reduce Aβ burden.

TAU tangles are another hallmark of AD.^57,58^ To further investigate the potential benefits of VIS, we employed a second *Drosophila* model of AD, in which the longest isoform of human TAU (2N4R) was expressed ubiquitously using *da*-GS and RU486.^59,60^ We employed the same schedule of RU486 feeding and vibration as before. We found that TAU expression significantly decreased sleep under no-vibration conditions (Figure 4, A-C). Importantly, vibration induced sleep in flies expressing TAU, demonstrating that vibration can effectively promote sleep in a second *Drosophila* model of AD. Furthermore, brain immunostaining using a TAU antibody revealed that vibration led to a profound decrease in TAU levels (Figure 4, D-E). Together, these results demonstrate that vibration promotes sleep in *Drosophila* models of AD and ameliorates the accumulation of Aβ and TAU.

**Figure 4.**
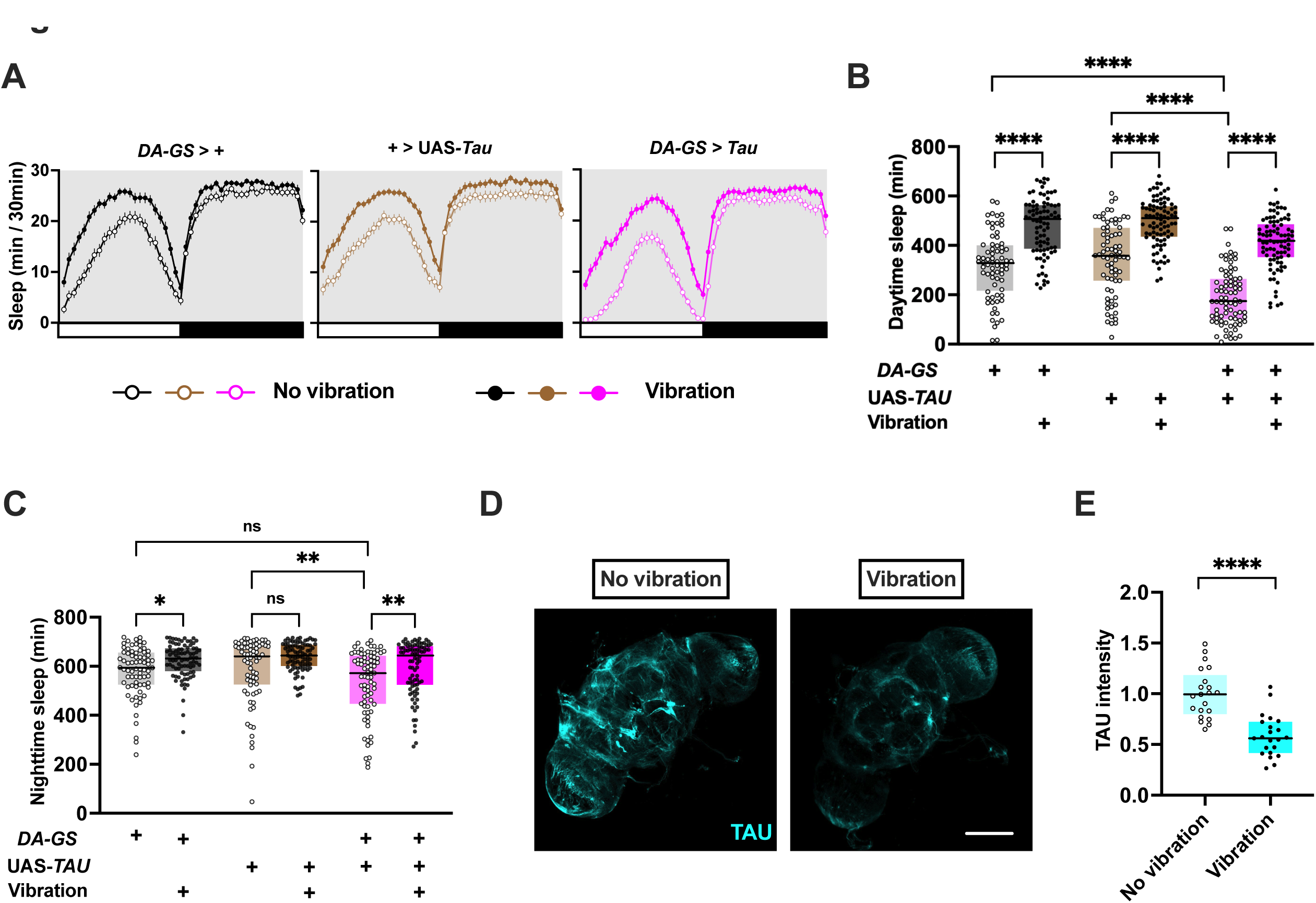
Vibration induces sleep in TAU-overexpressing flies and reduces TAU accumulation. **(A)** Sleep patterns of female flies from the specified genotypes and vibration conditions. Flies were exposed to continuous vibration for 3 days starting at ZT 0. The sleep amount for each 30- min bin was averaged over 3 days. Mean and SEM are shown. N =73-86. **(B-C)** Daytime (B) and nighttime (C) sleep for data shown in A. **(D)** Representative confocal images of immunolabeled TAU aggregates in the whole fly brains. Scale bar: 50 μm. **(E)** Quantification of TAU signal intensity normalized to the average intensity of the no-vibration condition. The line within the box denotes the median, while the box illustrates the 25^th^ to 75^th^ percentiles (B, C, E). Kruskal– Wallis test followed by Dunn’s multiple comparisons test (B, C); Mann-Whitney *U* test (E, F).

## Discussion

Our investigation demonstrates that sleep induced by gentle mechanical stimulation provides several cognitive and health benefits afforded by natural, uninduced sleep in *Drosophila*. VIS improved performance in a courtship conditioning paradigm, enhanced synaptic downscaling, and suppressed the accumulation of Aβ and TAU. These results are consistent with previous findings showing similar benefits of natural, uninduced sleep as well as genetically and pharmacologically induced sleep.^39,61,62^ These results, together with our previous findings showing the accumulation of sleep credit and increased arousal threshold during vibration, indicate that sleep induced by vibration and natural sleep share several key features and functions.

A recent study in zebrafish demonstrated that sleep-dependent synaptic downscaling was modulated by sleep pressure.^27^ It showed that synapse loss is most pronounced during sleep after prolonged wakefulness, which resulted in high sleep pressure. In contrast, pharmacologically induced sleep during periods of low sleep pressure failed to promote synaptic downscaling. These results suggest that the beneficial effects of VIS on synaptic downscaling may also depend on sleep pressure. More generally, VIS may enhance cognitive performance and clearance of toxic molecules only during periods of high sleep pressure. In our study, VIS likely occurred during periods of heightened sleep pressure, due to the activation of wake-promoting neurons or the expression of AD-associated molecules. Vibration administered during low sleep pressure may result in minimal cognitive and health benefits in *Drosophila*. Similarly, sleep induced by mechanosensory stimulation could be particularly beneficial to people experiencing sleep disruptions due to conditions like insomnia and neurodegenerative disorders but might not be as beneficial for healthy sleepers. The distinction could explain the varied findings on the effects of gentle rocking on cognition in humans. For instance, whereas one study found that rocking boosts sleep maintenance and memory in healthy sleepers,^6^ another study did not find significant effects of rocking on slow wave sleep and memory performance.^63^

Distinct sleep stages, such as slow wave sleep and rapid eye movement sleep, have long been recognized in mammals.^64^ Growing evidence suggests that *Drosophila* sleep also consists of multiple stages. Flies exhibit a greater increase in arousal threshold and a greater decrease in metabolic rate after prolonged sleep compared to the early stage of sleep, suggesting a transition from wakefulness to deep sleep through light sleep.^65,66^ Moreover, a recent study identified a discrete deep sleep stage where flies engage in proboscis extension to facilitate waste clearance, reminiscent of the glymphatic system in mammals.^67^ Our finding that VIS can enhance the clearance of Aβ and TAU suggests that vibration likely promotes the proboscis extension sleep stage. Additionally, although vibration provided only a partial rescue of the sleep loss due to the activation of wake-promoting neurons, it nearly completely rescued learning, memory and synaptic downscaling (Figures 1-2). These findings suggest that vibration may induce deep sleep under our experimental conditions. Further studies are required to determine how vibration influences different stages of sleep.

Research demonstrating the close link between sleep and neurodegenerative diseases suggests that sleep could help treat complex disorders such as AD. Our data demonstrate that VIS reduces the accumulation of AD-associated molecules, Aβ and TAU. Investigating whether VIS can also alleviate cognitive deficits linked to Aβ and TAU expression is an important future research direction. Although our primary focus was on the potential benefits of VIS in AD, it may hold promise for other neurodegenerative diseases associated with sleep disorders, such as Parkinson’s disease. However, since some *Drosophila* sleep mutants do not exhibit VIS,^7^ the first step should be to determine whether mechanosensory stimulation can improve sleep in various neurodegenerative diseases.

Previous studies illustrating cognitive and health benefits of sleep in animal models utilized pharmacological or genetic sleep induction methods.^68^ However, pharmacological sleep aids often induce considerable side effects due to their non-specific targeting, and genetic approaches to sleep induction are not currently feasible for human application. In contrast, anecdotal and experimental evidence supporting the sleep-promoting effects of rocking in humans suggests that gentle mechanosensory stimulation could offer a safer and more accessible option for sleep induction.^5,63^ This non-invasive approach warrants further investigation as a potential treatment option for patients with sleep disorders and neurodegenerative diseases.

## Acknowledgments

We thank the Bloomington *Drosophila* Stock Center, Janelia Fly Facility, Amita Seghal, and Mark Wu for fly stocks and William Joiner for SleepLab. This work was supported by a postdoctoral fellowship from Japanese Society for Promotion of Science (24KJ0033 to S.I.), grants from the National Institutes of Health (R21NS130878 and R01NS109151 to K.K.), and funds from Thomas Jefferson University Synaptic Biology Center (to K.K.).

## Data Availability

DAM locomotor data, video recordings, and confocal images reported in this article are available from the corresponding author upon request.

